## Supplementary Information for "Structural basis for peroxidase encapsulation in a protein nanocompartment"

### 1. Additional biophysical characterization of DyP-loaded KpEnc

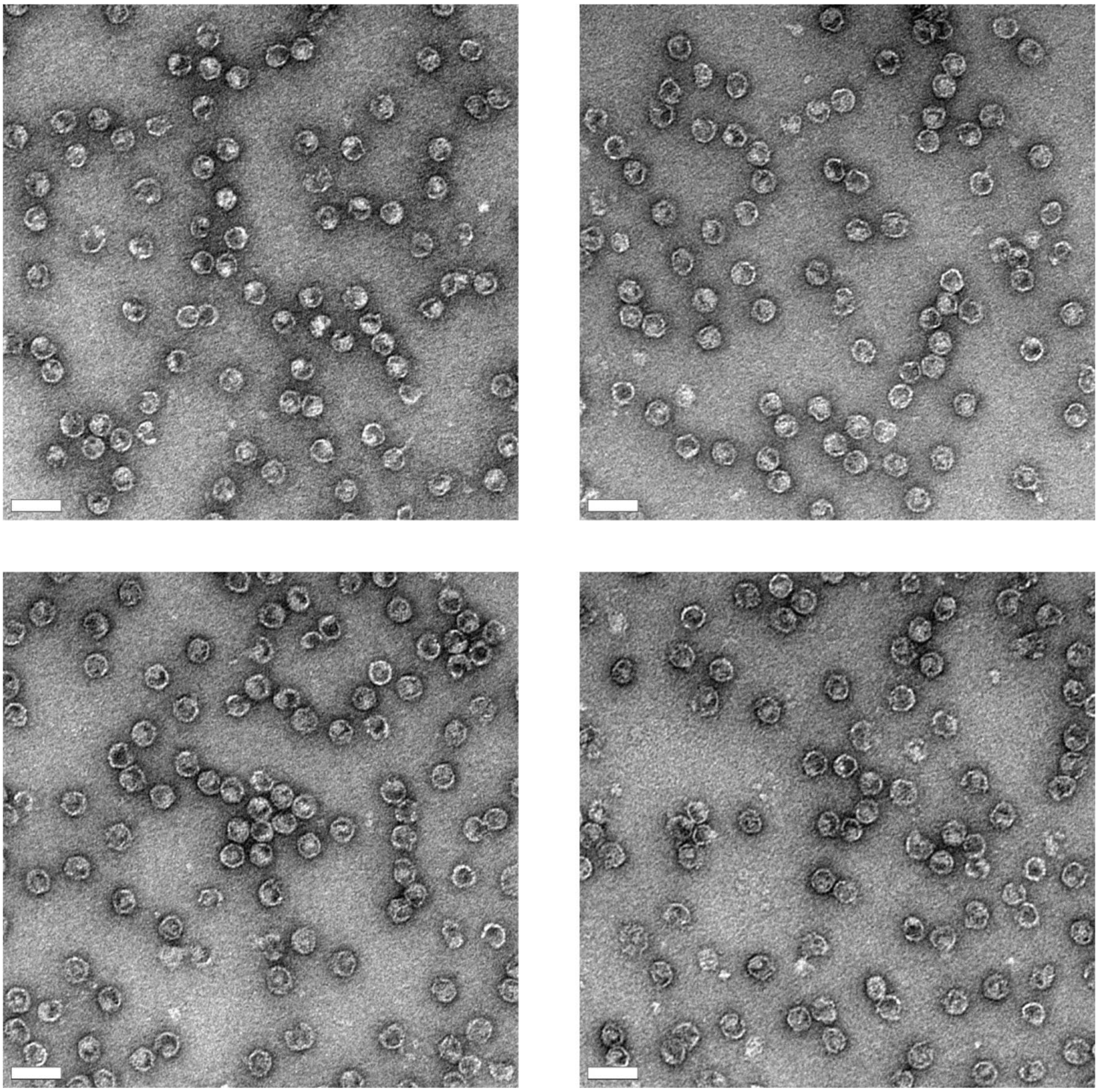

**Supplementary Fig. 1 | Additional TEM micrographs of DyP-loaded KpEnc nanocompartments.** TEM micrographs of separate KpDyP_Enc replicate samples. Scale bars (white): 50 nm.

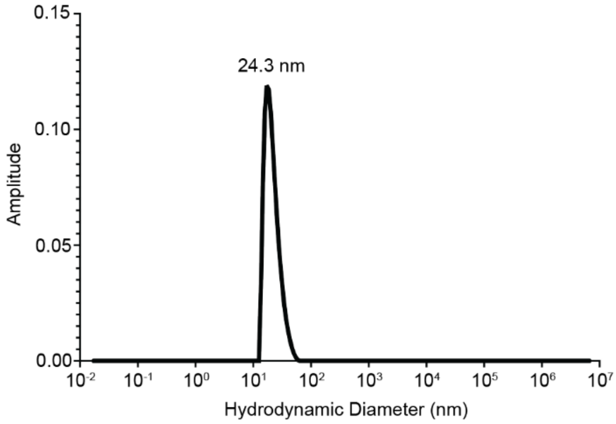

**Supplementary Fig. 2 | Dynamic light scattering analysis of DyP-loaded KpEnc.** DLS of DyP-containing KpEnc at pH 7.5 showing a Z-average diameter of 24.80 nm and a peak diameter of 24.30 nm.

### 2. Additional biophysical characterization of free KpDyP

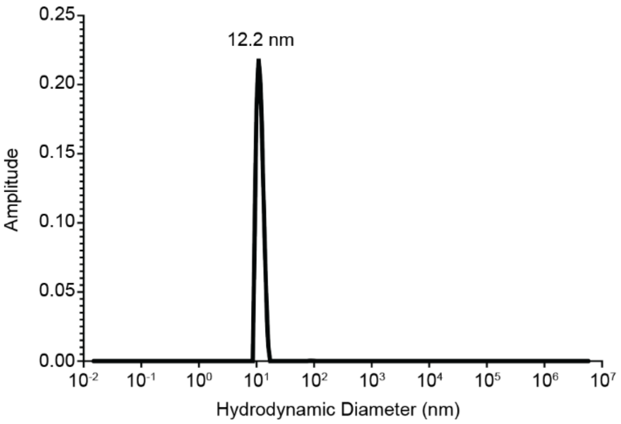

**Supplementary Fig. 3 | Dynamic light scattering analysis of free KpDyP.** DLS showing a Z-average diameter of 13 nm and a peak diameter of 12.21 nm for the purified free KpDyP at pH 7.5.

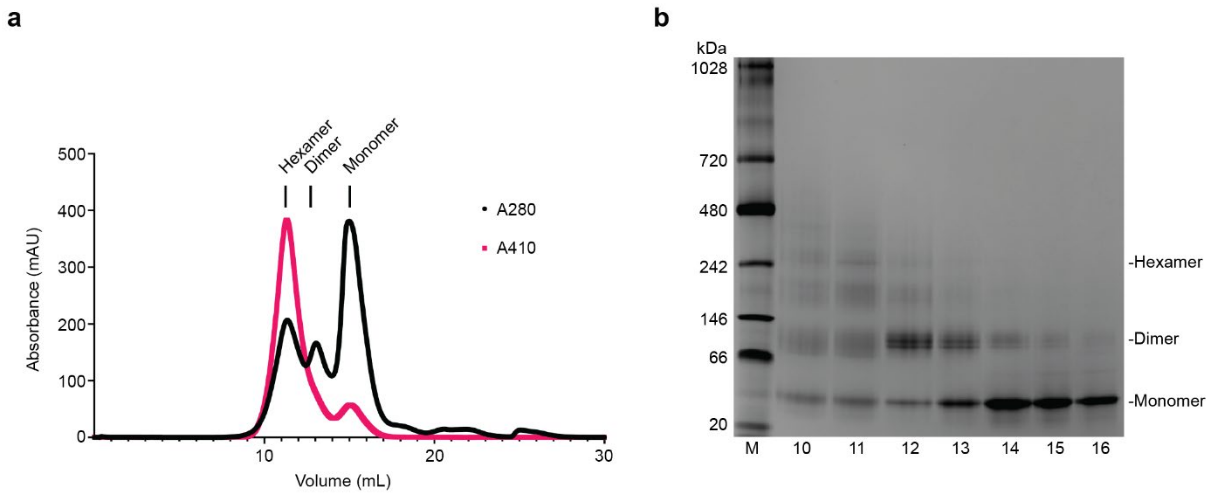

**Supplementary Fig. 4 | Analytical SEC and native PAGE analysis of free KpDyP.** **a,** Analytical SEC analysis of purified KpDyP using a Superdex 200 column. **b**, Native PAGE analysis of KpDyP-containing Superdex 200 fractions. Lane labels correspond to SEC mL. M, marker.

### 3. Cryo-EM analyses of free KpDyP

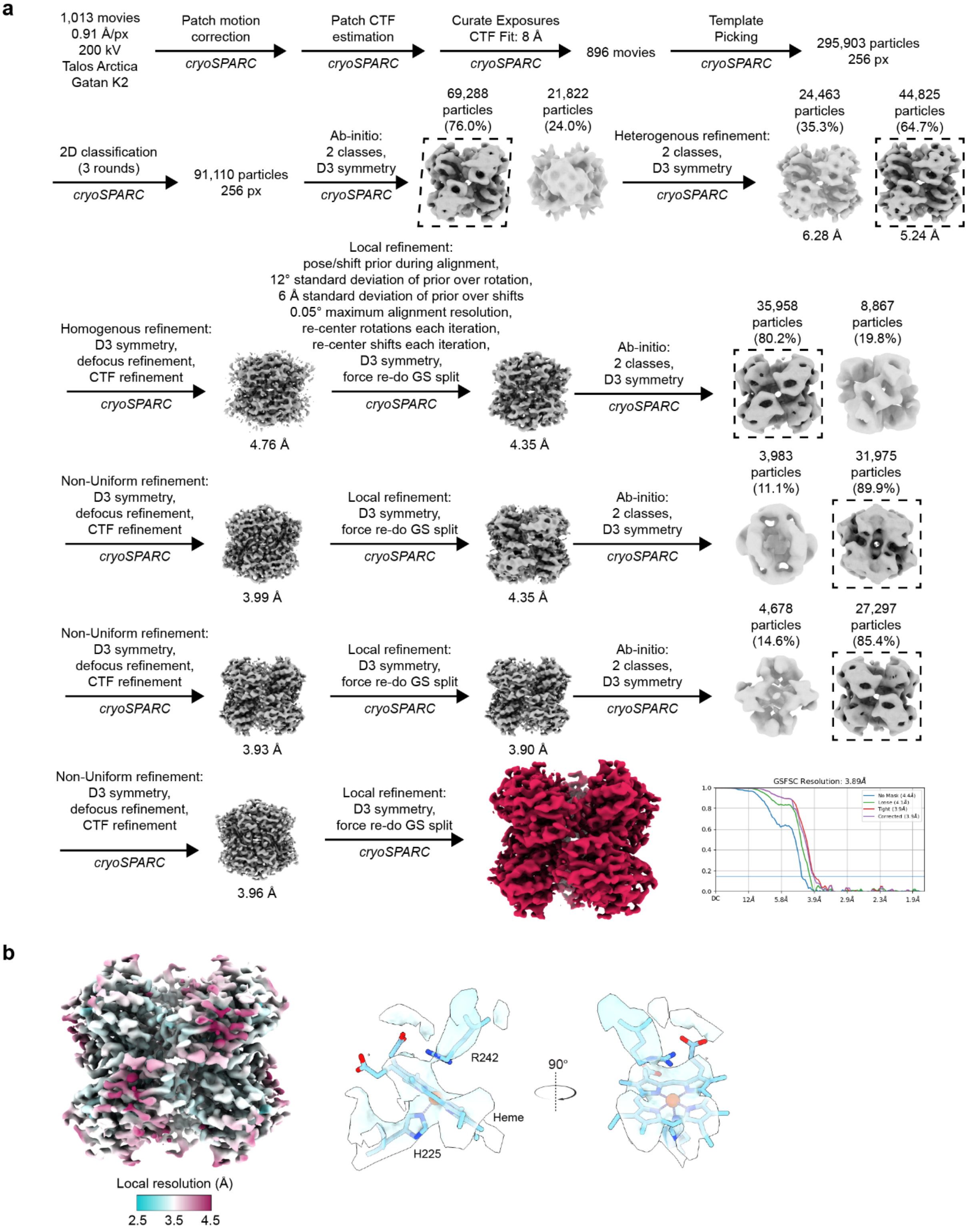

**Supplementary Fig. 5 | Cryo-EM analysis of free KpDyP.**[1,2] **a,** Cryo-EM data processing workflow. **b,** KpDyP hexamer colored by estimated local resolution (left) with magnified views of the heme density with atomic models for the heme moiety and His225 as well as Arg242 shown (middle and right).

**Supplementary Table 1 | Cryo-EM data collection and model building statistics.**[1,3-7]

| | KpDyP_Enc (EMD-41905) (PDB 8U50) | SUMO-TP_Enc (EMD-41906) (PDB 8U51) | KpDyP (EMDB-41904) (PDB 8U4Z) |
|---|---|---|---|
| **Data collection and processing** | | | |
| Magnification | 45,000x | 45,000x | 45,000x |
| Voltage (kV) | 200 | 200 | 200 |
| Electron exposure ($e^-/Å^2$) | 41.67 | 39.58 | 37.95 |
| Defocus range (μm) | -0.8 to -1.8 | -0.8 to -1.8 | -1.0 to -1.8 |
| Pixel size (Å) | 0.91 | 0.91 | 0.91 |
| Symmetry imposed | I | I | D3 |
| Initial particle images (no.) | 75,967 | 115,682 | 295,903 |
| Final particle images (no.) | 64,969 | 101,111 | 27,297 |
| Map resolution (Å) | 2.52 | 2.41 | 3.89 |
| FSC threshold | 0.143 | 0.143 | 0.143 |
| **Refinement** | | | |
| Initial model used (PDB code) | 7BOJ | 8U50 | AlphaFill |
| Model resolution (Å) | 2.8 | 2.7 | 4.4 |
| FSC threshold | 0.5 | 0.5 | 0.5 |
| Map sharpening *B* factor ($Å^2$) | -95.2 | -93.8 | -165.8 |
| Model composition | | | |
| Non-hydrogen atoms | 2,026 | 2,093 | 2,417 |
| Protein residues | 267 | 277 | 306 |
| Ligands | 0 | 0 | 1 |
| *B* factors ($Å^2$) | | | |
| Protein | 43.67 | 32.62 | 72.76 |
| Ligands | - | - | 67.82 |
| r.m.s. deviations | | | |
| Bond lengths (Å) | 0.004 | 0.005 | 0.004 |
| Bond angles (°) | 0.979 | 1.020 | 0.884 |
| Validation | | | |
| MolProbity score | 1.37 | 1.36 | 1.30 |
| Clashscore | 6.73 | 4.34 | 4.62 |
| Poor rotamers (%) | 0.47 | 1.36 | 0 |
| Ramachandran plot | | | |
| Favored (%) | 98.49 | 97.80 | 97.70 |
| Allowed (%) | 1.51 | 2.20 | 2.30 |
| Disallowed (%) | 0 | 0 | 0 |

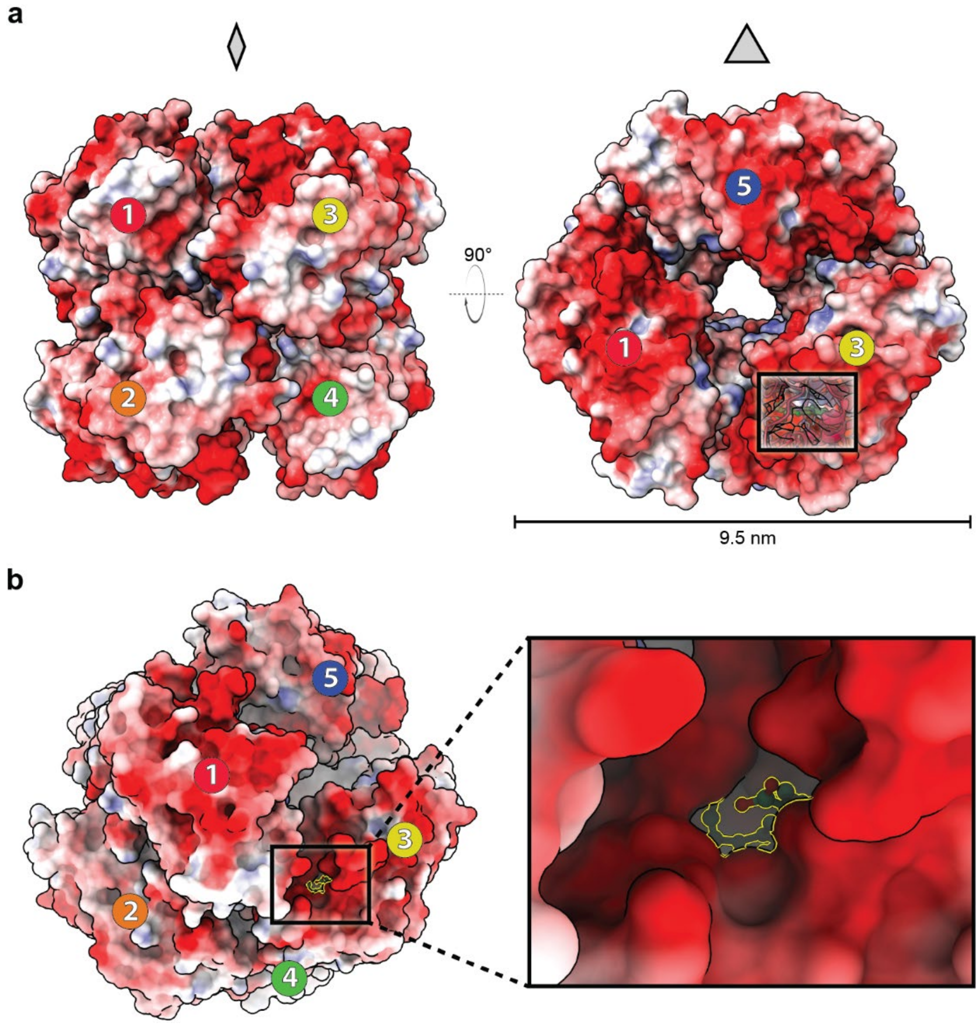

**Supplementary Fig. 6 | Electrostatic surface and active site entrance analysis of KpDyP.**[2] **a,** Electrostatic surface representation of the KpDyP hexamer along the two-fold symmetry axis (left) and three-fold symmetry axis (right) with individual subunits numbered and heme molecule highlighted (yellow) through transparent surface (right, black box). Theoretical isoelectric point (pI) was calculated to be 4.65 and the charge at pH 7.0 was calculated to be -21.70, with 54 negatively charged residues and 32 positively charged residues. **b,** Rotated model showing heme (highlighted yellow) as well as a zoomed in representation of a potential access channel to the active site (right).

### 4. Cryo-EM analysis of DyP-loaded KpEnc (KpDyP_Enc)

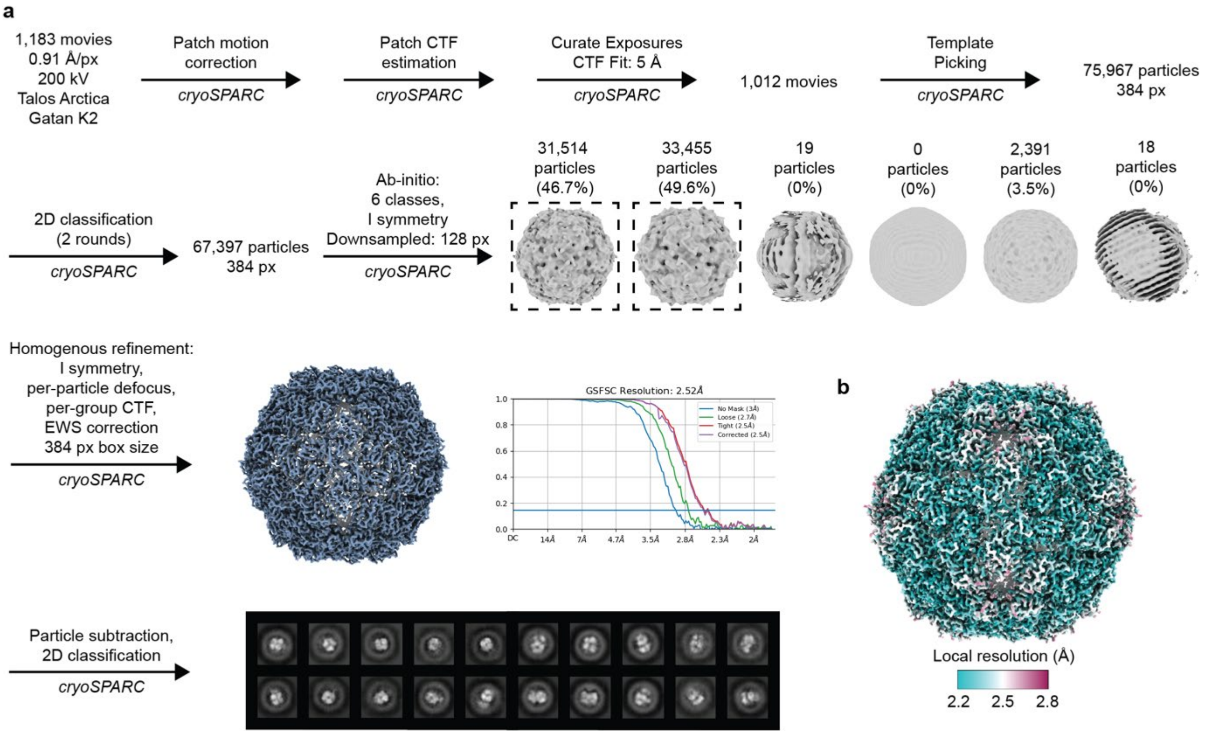

**Supplementary Fig. 7 | Cryo-EM analysis of KpDyP_Enc.**[1,2] **a,** Cryo-EM data processing workflow. **b,** Exterior view of the KpDyP_Enc shell colored by estimated local resolution.

### 5. KpEnc pore analysis

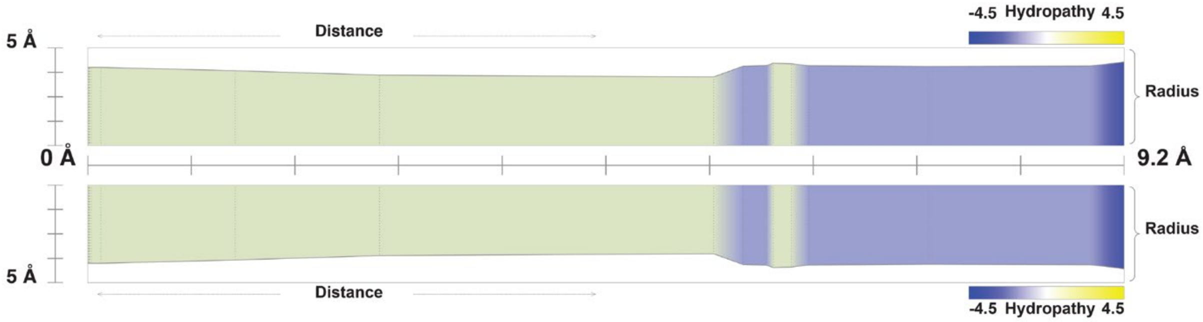

**Supplementary Fig. 8 | KpEnc 5-fold pore characterization via MOLEonline.**[8,9]

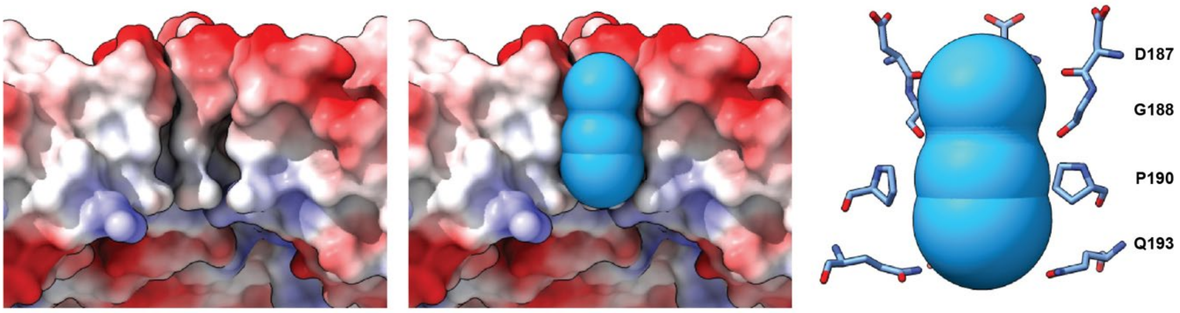

**Supplementary Fig. 9 | Structural analysis of the KpEnc 5-fold pore using MOLEonline.**[8,9] Electrostatic surface representation of the 5-fold pore. The residues of one subunit lining the 5-fold pore are shown.

### 6. Cryo-EM analysis of SUMO-loaded KpEnc (SUMO-TP_Enc)

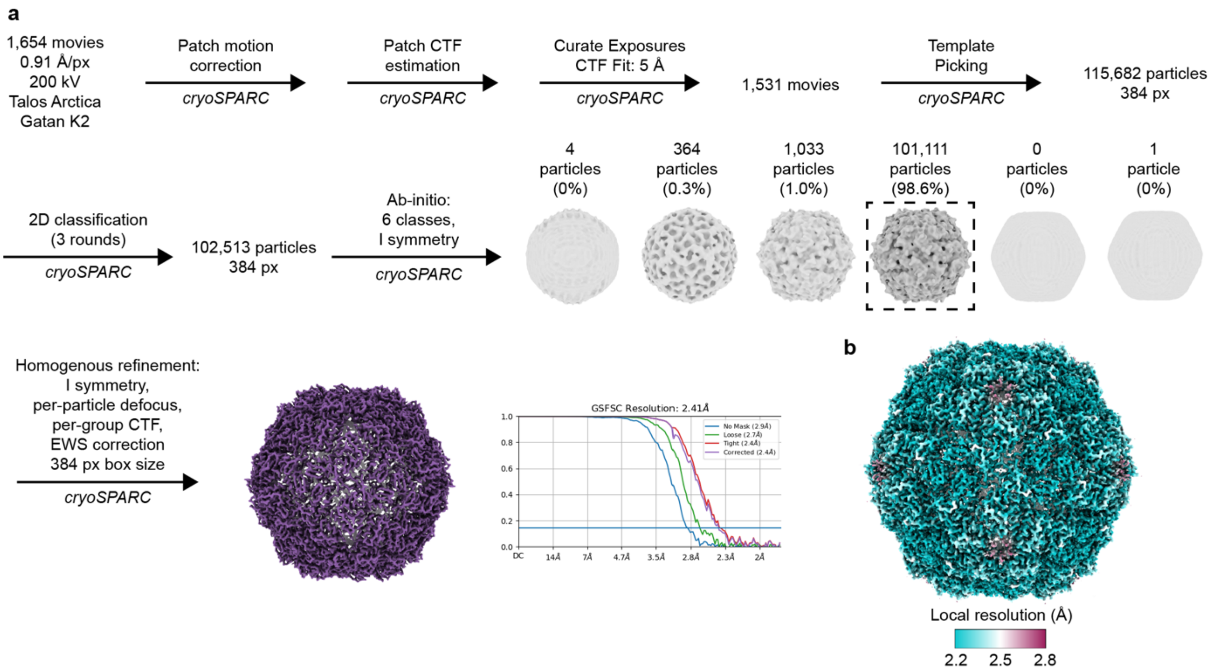

**Supplementary Fig. 10 | Cryo-EM analysis of SUMO-TP_Enc.**[1] **a,** Cryo-EM data processing workflow. **b,** Exterior view of the SUMO-TP_Enc shell colored by estimated local resolution.

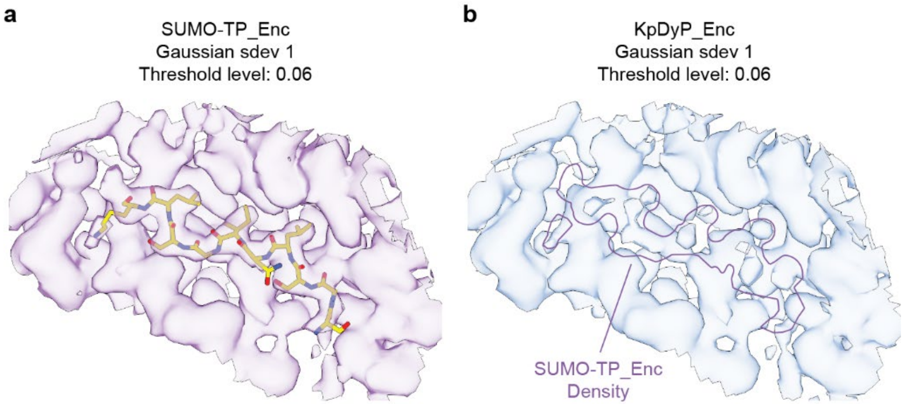

**Supplementary Fig. 11 | Comparison of the TP binding site densities of KpDyP_Enc and SUMO-TP_Enc.**[1,2] **a,** Cryo-EM density map of the KpEnc encapsulin interior surface and the bound TP from SUMO-TP_Enc (purple) with SGSLNIGSLK TP in stick representation (yellow) for map-to-model comparison. **b,** Cryo-EM density map of the TP binding site of KpDyP_Enc (blue). The outline of the TP density from SUMO-TP_Enc is shown and overlayed for comparison (purple).

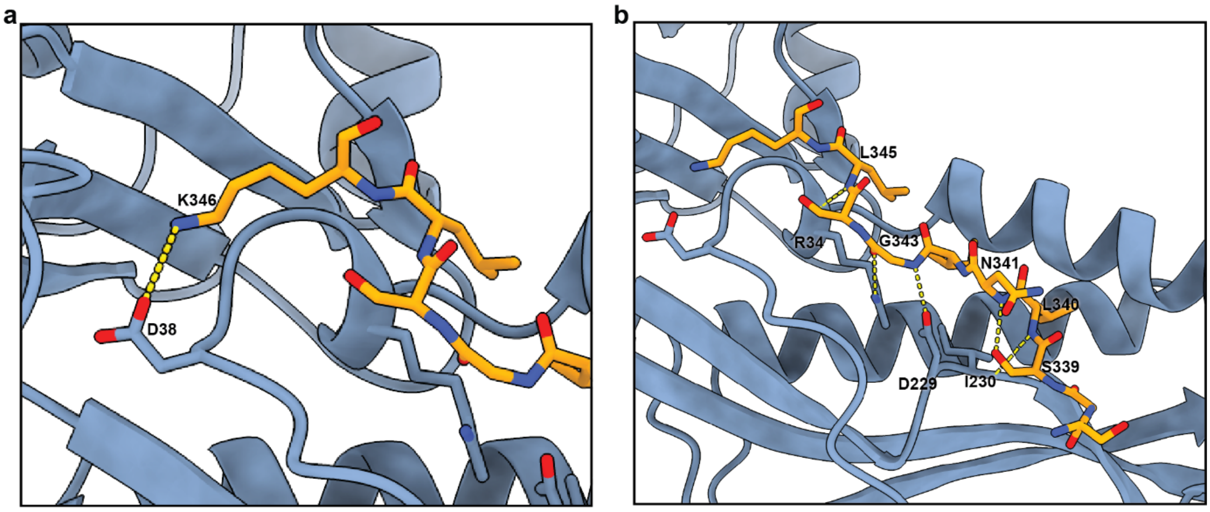

**Supplementary Fig. 12 | Ionic and hydrogen bonding interactions mediating TP-shell binding. a**, Salt bridge between TP residue Lys 346 and encapsulin shell protein residue Asp38. **b**, Inter- and intramolecular hydrogen bonds of TP residues.

### 7. SDS-PAGE gel TP mutant purifications

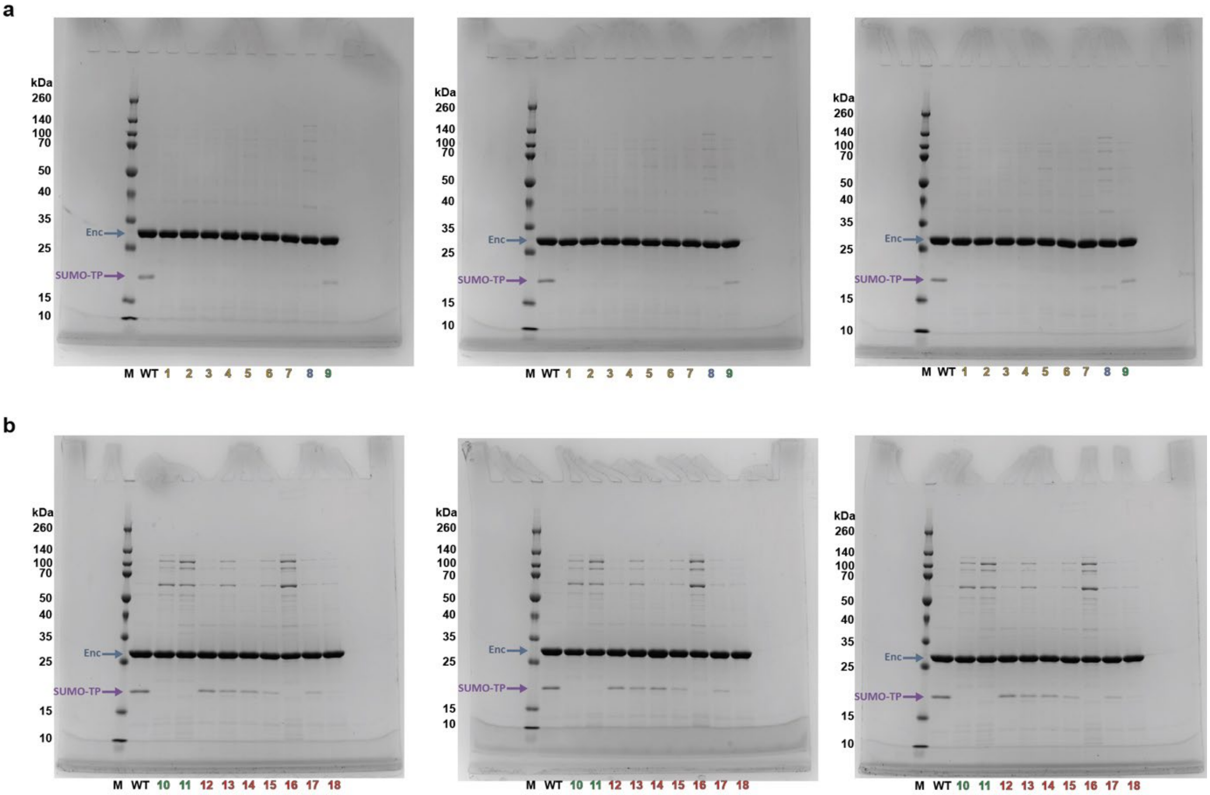

**Supplementary Fig. 13 | Full SUMO-TP mutant SDS-PAGE gels. a,** Full triplicate SDS-PAGE gels illustrating cargo loading capacity of WT and respective mutants 1 through 9 with KpEnc (higher band, blue) and the co-purified encapsulated SUMO-TP mutant cargo (lower band, purple) where present. **b**, Same layout as above, but for mutants 10 through 18.

### 8. Protein sequence data

**Supplementary Table 2 | Protein sequences of constructs used in this study.**

| Construct | Protein sequence[1,2] |
|---|---|
| KpDyP_Enc | MACPISQSVSQPVDERLTRAAIFLVVTINPGKAAEVAVRTLCGTLSSLVR<br>GVGFRILDGGLSCVMGVSSGGWERLFGDEKPEYLHVFQEINGVHHAPSTP<br>GDLLFHIRAARMDLCFELASRILSDLGSSVCVVDSVQGFRYFDDRDLLGF<br>VDGTENPVAQAAVDATLIGEEDHTFSGGSYVIVQKYLHDLDKWNAIPVEQ<br>QEKIIGREKLSDIELKDADKPSYAHNVLTSIEEDGEDVDILRDNMPFGDP<br>GKGEFGTYFIGYSRKPARIERMLENMFVGNPPGNYDRILDVSRAITGTLF<br>FIPTVSFLDSVEPQPSVSQQTDDAKYIYDSPGTKGESGSLNIGSLKKEVQ<br>DE*…<br>MNNLHRELAPVSDAAWEQIEEEASRTLKRFLAARRVVDVSDPQGPA<br>FSAVGTGHVTRLEGPGDSVGAVKRQSQPVVEFRVPFILTRQAIDDVERGS<br>QDSDWSPLKEAARKIAGAEDRAVFDGYAAAGIGGIRPQSSNSPLTLPVAA<br>SGYPDVIARALDQLRVAGVNGPYHLVLGENAYTLITSGNEDGYPVLQHIH<br>RLIDGEIVWAPAIEGGVLLSTRGGDFAMDIGQDISIGYLSHTATHVELYL<br>QESFTFRTLTSEAVVSLLPSED* |
| KpDyP | MACPISQSVSQPVDERLTRAAIFLVVTINPGKAAEVAVRTLCGTLSSLVR<br>GVGFRILDGGLSCVMGVSSGGWERLFGDEKPEYLHVFQEINGVHHAPSTP<br>GDLLFHIRAARMDLCFELASRILSDLGSSVCVVDSVQGFRYFDDRDLLGF<br>VDGTENPVAQAAVDATLIGEEDHTFSGGSYVIVQKYLHDLDKWNAIPVEQ<br>QEKIIGREKLSDIELKDADKPSYAHNVLTSIEEDGEDVDILRDNMPFGDP<br>GKGEFGTYFIGYSRKPARIERMLENMFVGNPPGNYDRILDVSRAITGTLF<br>FIPTVSFLDSVEPQPSVSQQTDDAKYIYDSPGTKGESGSLNIGSLKKEVQ<br>DEENLYFQGGSGGHHHHHH* |
| SUMO-TP_KpEnc | MSDSEVNQEAKPEVKPEVKPETHINLKVSDGSSEIFFKIKKTTPLRRLME<br>AFAKRQGKEMDSLRFLYDGIRIQADQTPEDLDMEDNDIIEAHREQIGGGG<br>SGGSGGSGGPGTKGESGSLNIGSLKKEVQDE*… MNNLHRELAPVSDAA<br>WEQIEEEASRTLKRFLAARRVVDVSDPQGPAFSAVGTGHVTRLEGPGDSV<br>GAVKRQSQPVVEFRVPFILTRQAIDDVERGSQDSDWSPLKEAARKIAGAE<br>DRAVFDGYAAAGIGGIRPQSSNSPLTLPVAASGYPDVIARALDQLRVAGV<br>NGPYHLVLGENAYTLITSGNEDGYPVLQHIHRLIDGEIVWAPAIEGGVLL<br>STRGGDFAMDIGQDISIGYLSHTATHVELYLQESFTFRTLTSEAVVSLLP<br>SED* |

[1] Intergenic sequences for encapsulated constructs represented by ellipses.
[2] Stops indicated by asterisks.
